# Mitigating mitochondrial genome erosion without recombination

**DOI:** 10.1101/143586

**Authors:** Arunas L Radzvilavicius, Hanna Kokko, Joshua Christie

## Abstract

Mitochondria are ATP-producing organelles of bacterial ancestry that played a key role in the origin and early evolution of complex eukaryotic cells. Most modern eukaryotes transmit mitochondrial genes uniparentally, often without recombination among genetically divergent organelles. While this asymmetric inheritance maintains the efficacy of purifying selection at the level of the cell, the absence of recombination could also make the genome susceptible to Muller’s ratchet. How mitochondria escape this irreversible defect accumulation is a fundamental unsolved question. Occasional paternal leakage could in principle promote recombination, but it would also compromise the purifying-selection benefits of uniparental inheritance. We assess this tradeoff using a stochastic population-genetic model. In the absence of recombination, uniparental inheritance of freely segregating genomes mitigates mutational erosion, while paternal leakage exacerbates the ratchet effect. Mitochondrial fusion-fission cycles ensure independent genome segregation, improving purifying selection. Paternal leakage provides opportunity for recombination to slow down the mutation accumulation, but always at a cost of increased steady-state mutation load. Our findings indicate that random segregation of mitochondrial genomes under uniparental inheritance can effectively combat the mutational meltdown, and that homologous recombination under paternal leakage might not be needed.

## Introduction

Mitochondria are descendants of free-living bacteria that became endosymbiotic within an archaeal host cell at the dawn of eukaryote evolution (Martin et al., 2015). Most of the proto-mitochondrial endosymbiont genes were either lost or transferred to the nucleus, leaving a diminutive genome of 37 genes in vertebrates, and up to around 100 genes in early-branching eukaryotes (Lang et al., 2003). Oxidative phosphorylation (OXPHOS)—the most critical function of mitochondria in modern eukaryotes—depends on the genomic stability and maintenance of these genes, as well as interactions between mitochondrial and nuclear genes that both encode subunits of respiratory-chain protein complexes. Mitochondrial mutations result in debilitating diseases and neuromuscular deterioration in humans (Taylor and Turnbull, 2005; Wallace, 2010), while mitochondrial-nuclear mismatches have been shown to induce negative developmental, fertility and cognitive effects in laboratory animal studies (Wolff et al., 2014).

Eukaryotic sex involves inheritance of nuclear genes from both parents, but is highly asymmetric in transmission of mitochondrial genes that in higher eukaryotes are predominantly inherited from the maternal gamete. This asexual mode of mitochondrial transmission, along with reduced effective population size and relatively high nucleotide substitution rates, has been suggested to cause gradual deterioration of the mitochondrial genome (Lynch, 1996; Lynch et al., 2006; Neiman and Taylor, 2009; Greiner et al., 2014) through recurrent stochastic losses of the least-loaded genome class—a concept known as Muller’s ratchet (Muller, 1964; Felsenstein, 1974). Muller’s ratchet in asexual endosymbiont genomes was likely one of the major forces driving an early massive gene transfer form proto-mitochondria to the emerging eukaryotic nucleus (Martin and Hermann, 1998; Timmis et al., 2004), establishing the nuclear-mitochondrial asymmetry in genome size. Nevertheless, a handful of essential genes remain localized within modern mitochondria (Race et al., 1999), and were lost only in mitochondrion-derived organelles that do not perform oxidative phosphorylation, such as hydrogenosomes and mitosomes. There is therefore a strong evolutionary pressure for retention of these genomic outposts at the energy-generating membranes, which requires mechanisms mitigating mutational deterioration.

Multiple lines of empirical evidence suggest that mitochondrial genomes of modern eukaryotes could be protected against a ratchet-like mutational meltdown (Rand, 2008; Stewart et al., 2008). Unlike the mammalian Y chromosome, the animal mitochondrial genome is not subject to the accumulation of transposable elements, and has a remarkably stable gene content (Boore, 1999). Additionally, non-synonymous substitution rates for mitochondrial genes coding for core respiratory subunits are in many cases comparable to or lower than substitution rates in nuclear loci (Zhang and Broughton, 2013; Popadin et al., 2013; Cooper et al., 2015), and free-living prokaryotes (Itoh et al., 2002). This implies strong purifying selection against mitochondrial mutations, and challenges the conventional prediction that non-recombining animal mitochondrial genomes are subject to excessive accumulation of detrimental substitutions (Lynch, 1996; Lynch and Blanchard, 1998; Neiman and Taylor, 2009).

Uniparental inheritance of mitochondrial genes (UPI) facilitates purifying selection at the level of the cell by maintaining high cell-to-cell variance in mutation load (Bergstrom and Pritchard, 1998; Hadjivasiliou et al., 2013; Christie and Beekman, 2017). Furthermore, UPI has important implications for heteroplasmy (Christie et al., 2015), adaptive evolution (Christie and Beekman, 2017) and mito-nuclear coadaptation (Hadjivasiliou et al., 2012; 2013), and could have played an important role in the origin of self-incompatible mating types and evolution of sexual dimorphism in higher metazoans (Hurst and Hamilton, 1992; Radzvilavicius et al., 2016). The rule of strict UPI can be partially broken (termed paternal leakage) or completely absent (biparental inheritance). Under those conditions, theoretical modelling makes opposite predictions: less efficient selection against defective cytoplasmic genes, increased mutational load at equilibrium and easier spread of selfish genetic elements (Roze et al., 2005; Bastiaans et al., 2014).

These theoretical arguments suggest that asymmetric inheritance plays an important role in keeping mitochondria healthy, but it is not clear whether purifying cell-level selection alone can provide sufficient protection against Muller’s ratchet in small populations. UPI promotes mitochondrial clonality (homoplasmy) within the cell, and therefore limits the scope and potential effects of homologous recombination, and without recombination the population-wide fixation of mutations is irreversible. It has been proposed that the long-term stability of mitochondrial genome requires episodic reversion to biparental transmission (paternal leakage), which elevates effective recombination rates and is thus argued to slow down mitochondrial genome erosion (Hoekstra, 2000; Neiman and Taylor, 2009; Dokianakis and Ladoukakis, 2014; Greiner et al., 2014).

While homologous recombination is the key mechanism countering Muller’s ratchet under haploid population genetics (Felsenstein, 1974), the interplay between mutation accumulation and recombination in organelle genomes is more complex and less well understood. Mitochondrial DNA exists in a nested hierarchy of several genome copies within a mitochondrial nucleoid (Satoh and Kuroiwa, 1991; Jacobs et al., 2000), dozens of nucleoids within an organelle and multiple organelles per cell (Satoh and Kuroiwa, 1991). Selection against deleterious mitochondrial mutations therefore operates mostly through their effects on the host cell fitness, that is, at the level of the group of mitochondrial genomes. The composition of these genome groups changes due to random organelle segregation at cell division, stochastic sampling in bottleneck-like processes, paternal leakage and recombination. Cell-level (and organism-level) performance may not be strongly compromised if only one or a few of its many mitochondria acquire mutations (Rossignol et al., 2003).

Random mitochondrial segregation at cell division increases mutational variance, meaning that mitochondrial mutation load of the daughter cell could markedly differ from the parent. Relative to strict uniparental transmission, paternal leakage reduces this variance, which hinders the host-level selection against deleterious mutations and increases steady-state mutation load (Bergstrom and Pritchard, 1998; Hadjivasiliou et al., 2013). But without paternal leakage, homologous recombination has little effect, as intra-cellular variance in this case comes only from de-novo mutations. There is therefore a tradeoff between the two mechanisms: paternal leakage increases the opportunity for mitochondrial recombination, but reduces the efficacy of selection at the level of the host cell. To the best of our knowledge, there is no formal theory examining how the balance between uniparental inheritance, on the one hand, and paternal leakage and recombination, on the other, affects the accumulation of deleterious mutations.

To understand the dynamics of mitochondrial mutation accumulation under segregational drift, paternal leakage and homologous recombination, we developed a population genetic model of unicellular eukaryotic species subject to purifying host-level selection. Consistent with previous studies, we find that paternal leakage relaxes selection against defective mitochondrial genes, and in the absence of recombination severely increases the rate of mutation fixation. Clustering of mitochondrial DNA into strongly linked groups (such as nucleoids or organelles), and large mitochondrial population sizes reduce segregational drift and further accelerate genome degradation. Strict uniparental inheritance and tight bottlenecks in mitochondrial population size, on the other hand, provide protection against Muller’s ratchet even without recombination, due to strong purifying selection and increased cell-to-cell variance in mutation load. When there is paternal leakage, homologous recombination can reduce the rate of mutation fixation, but the increase in steady-state mutational load due to mitochondrial mixing remains. Taken together, our results indicate that random segregational drift in uniparental inheritance alone could mitigate the mutational meltdown in mitochondrial genes, and that homologous recombination might not be necessary.

## Methods

We developed a stochastic model representing a finite population of unicellular eukaryotes, containing *M* mitochondrial genomes each. Each discrete generation consists of non-overlapping steps of (1) mitochondrial mutation, (2) mating with cytoplasmic mixing in the form of paternal leakage, (3) cell division with a bottleneck and (4) selection. This particular order of events is representative of a haploid life cycle (i.e. selection acts after syngamy and meiosis), but the order could be altered without major implications to our main conclusions. The population size is fixed at *N*. Definitions of symbols and model parameters are given in Table 1. The model was implemented in C++ with the source code available at *https://github.com/ArunasRadz.vilavicius/Mutations*.

### Mutation

The mitochondrial genome is modelled as a set of *K* loci, each locus representing a segment that can be replaced by a single recombination event. At the start of a generation, each genome within the cell acquires a Poisson-distributed number of new mutations (mean *μ*). These new mutations are distributed randomly among the *K* loci, independent of the current mutational state of a locus. We track the number of deleterious point mutations within each locus assuming that this has no upper limit and so ensuring that the pace of mutation fixation (d*m*/d*t*) under the multiplicative fitness function remains constant (Takeuchi et al., 2014). Back-mutations are ignored, so that the fixation of a mutant allele within a locus is irreversible.

### Paternal leakage and recombination

We consider two major levels of gene mixing: paternal leakage and recombination. Following mutation, we randomly divide the population into *N/2* pairs for mating, assuming no mating types or sexes. Each pair of cells exchanges a Poisson-distributed number of mitochondria (mean L). Since the number of organelles exchanged between mating partners cannot exceed *M,* we use a truncated Poisson distribution and consider only values of paternal leakage *L* between 0 and M/2. The number of mitochondrial genomes exchanged is drawn independently for each mating pair.

The next step is homologous recombination within the cell. This is modelled as the exchange of a Poisson-distributed number of alleles (mean *R* per cell) between mitochondrial genome pairs. The participating genomes are chosen randomly for each homologous gene transfer event, as is the recombining locus. Gene transfer between mitochondria is unidirectional and thus resembles horizontal gene transfer in prokaryotes (Takeuchi et al., 2014), which is likely a simplification of reality.

### Cell division

Each cell replicates its mitochondrial population by clonal doubling, after which *M* randomly chosen mitochondrial genomes are transmitted to a daughter cell (sampling without replacement). This process of random segregation increases cell-to-cell variance in mitochondrial mutational load, producing daughter cells that can carry more or fewer mutations than the parent. To model the clustering of mitochondrial genomes into nucleoids or organelles, we assume *M/C* clusters in which *C* mitochondrial genomes are tightly linked. At cell division, each cluster segregates as a unit. With *C*=1 (our default assumption) all *M* mitochondrial genomes replicate and segregate independently, while with *C*=*M* the daughter cell is a clonal copy of the parent. However, the genomic composition of the cluster is not necessarily permanent. We therefore consider random redistribution of mitochondrial genomes within the cell (between clusters, e.g. between mitochondria with multiple mtDNA molecules), which precedes cell division. This is modelled as an exchange of mitochondrial genomes within the same cell, with the total number of genome pairs that exchange their locations *F*_tot_ following the Poisson distribution (mean migration rate *F*). When *F*=0, mtDNA clusters are permanently linked, and replicate as a cohesive whole. With high values of F, mtDNA packaging into clusters is random, i.e. clusters are regenerated from the whole mtDNA population of the cell before each cell division.

Finally, we consider the effect of mitochondrial bottlenecks. Following cell division, the mitochondrial genome population is reduced through random sampling without replacement from *M* down to *B,* and then increased back to *M* through error-free replication. Lower values of *B* therefore represent tighter bottlenecks. The bottleneck is simply a mechanism of reducing mutational variance within the cell (increasing homoplasmy), and the precise details of how this is achieved in real biological systems (Cao et al., 2007, 2009; Wai et al., 2008; Johnston et al., 2015) are not relevant for our purposes.

### Selection

The life cycle ends with selection among cells, which we model as weighted random sampling of *N* individuals with replacement, with cell fitness values forming the weights. All mutations are assumed to contribute equally to the deleterious fitness effect without epistasis, so that the fitness contribution of a mitochondrial genome with *m* point mutations is *w_m_* = (1 − *s*)^*m*^. Cell fitness is then the arithmetic mean of its *M* mitochondrial fitness contributions. This does not account for the inter-mitochondrial epistatic interactions (Hadjivasiliou et al., 2013), but guarantees that d*m*/d*t* does not depend on the total mutational load of the cell. Surviving cells give start to a new generation.

Scaling normalization approaches derive their technical bias estimates from ratio of proportions.

**Table 1. List of parameters and symbols**
|  |  |
| --- | --- |
| $N$ | Population size |
| $\mu$ | Mutation rate per mitochondrial genome per generation |
| $s$ | Fitness effect of a mitochondrial mutation |
| $M$ | Number of mitochondrial genomes per cell |
| $L$ | Paternal leakage, mitochondria per cell per generation |
| $R$ | Mitochondrial recombination rate, per cell per generation |
| $K$ | Number of mitochondrial loci |
| $B$ | Bottleneck size |
| $C$ | Mitochondrial cluster size |
| $F$ | Genome migration rate within the cell |
| $H_{LLC(+1)}$ | Gene diversity of the least-loaded (second least-loaded) class |

## Results

### Dynamics of mitochondrial mutation accumulation and fixation

We first explored the general behavior of the model in the absence of mitochondrial recombination by following mutation accumulation and population-wide fixation in freely-segregating mitochondria (our default assumption of *C*=1, Fig. 1). We also analyzed the time-evolution of the average gene diversity of the least-loaded mitochondrial genome class *H*_LLC_ and that of the second-least loaded class *H*_LLC+1_, defined as 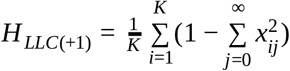, where *x_ij_* is the frequency of the allele with *j* point mutations at locus *i*.

**Figure 1.**
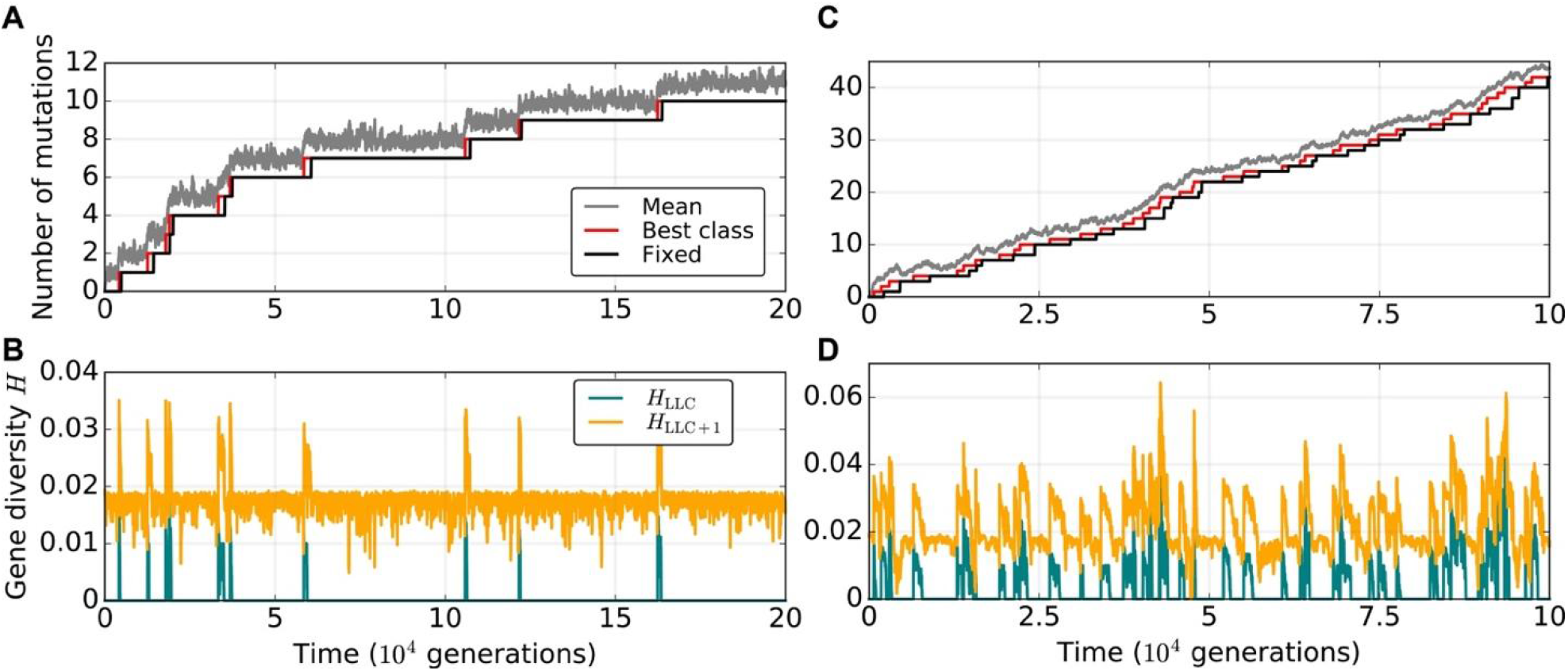
Mutation accumulation profiles in mitochondrial genomes of a small eukaryotic population with no recombination. (A, C) Population mean of the deleterious mutation load (gray), the number of mutations in the least-mutated genome class (red) and the number of mutations fixed within the population (black). (B, D) Gene diversity in the least-loaded (teal, *H*_LLC_) and the second least-loaded class (orange, *H*_LLC+1_). Population size is set to *N*=500, *M*=20, *μ*=0.005, C=1. Here and in the rest if the figures, *s*=0.02 and *K*=100. The rate of paternal leakage is *L*=0.8 (A, B) or *L*=5.0 (C, D).

Stochastic drift in the model population operates at two major levels—within the eukaryotic population of size *N* and through random segregation of mitochondrial haplotypes at cell division. The mitochondrial population distributed over *N* eukaryotic cells can be further subdivided into classes according to the number of deleterious mutations *m*. In populations of finite size, the genome class containing the fewest mutations *m* _LLC_ will eventually be lost because of stochasticity, and, in the absence of back mutations and recombination, won’t be recovered, causing the continuous accumulation of mutant alleles.

The analysis of mutation-accumul ation profiles recapitulates the main conclusions of Charlesworth and Charlesworth (1997) and Takeuchi et al. (2014), who studied haploid populations and found that each advance of Muller’s ratchet (i.e. each stochastic loss of the least-mutated genome class) is followed by fixation of a single deleterious mutant allele within the whole population (Fig. 1). In a quasi-steady state—that is, between the stochastic mutation fixation events—the least-loaded class shows no diversity (*H*_LLC_=0) while the value of *H*_LLC+1_ remains close to 0.02 for the number of mitochondrial loci K=100. This indicates that at equilibrium the second least loaded class is well approximated by *K* distinct genotypes of equal frequencies, each with one freely segregating mutation per locus 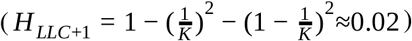. The moment the least-mutated class is lost, the value of *H*_LLC_ becomes equal to the former *H*_LLC+1_, but then rapidly returns to zero due to drift, indicating the fixation of a single mutant allele in the new least-loaded genome class. This is then followed by the drop of the new *H*_LLC+1_ back to its steady-state value, with one segregating mutation per locus (Fig. 1B). Since all genome classes containing more than *m*_LLC_ mutations are ultimately derived from the least-mutated class, the fixation of a mutation within the fittest genome is followed by fixation in the second least-loaded class, and the whole population.

### Uniparental inheritance mitigates mitochondrial mutation accumulation

We next systematically explored the effect of parameters controlling the strength of segregational drift and genetic mixing at both levels of organization. For each set of parameter values, we tracked the population evolution for at least 10^6^ generations, measuring the population mean of mitochondrial mutation load, the number of mutations in the least-mutated genome class, and the number of deleterious mutations fixed within the population.

Uniparental inheritance maintains highly efficient purifying selection against mitochondrial mutations in large part due to the segregational drift at each cell division, where each daughter cell inherits a random sample of the parental cell’s mitochondrial population. In other words, selection operates more efficiently through the recurrent creation of individual cells that are better or worse—with respect to deleterious mutations—than their parental cells. Previous studies have shown that mitochondrial mixing through biparental inheritance reduces variance in mutational load between lineages of eukaryotic cells with freely-segregating mitochondria (Hadjivasiliou et al., 2013; Radzvilavicius, 2016; Christie and Beekman, 2017). This reduced variance resulted in weaker purifying selection at the level of the cell (groups of mitochondria), and higher equilibrium mutation load in infinite eukaryotic populations. Here we find that without recombination, even moderate levels of paternal leakage *L* severely increase the rate of mutation accumulation (Fig. 1C, and Fig. 2), and could induce a dynamical regime in which the loss of the least-loaded genome class occurs several times before the corresponding number of mutations become fixed within the entire population (Fig. 1D). Strictly uniparental inheritance of mitochondria, on the other hand, is capable of maintaining negligible fixation rates of deleterious mutations (Fig. 2). Increasing the size of the mitochondrial population *M* reduces the effect of stochastic drift between cell divisions and results in lower variance in mutational load. This hinders purifying selection at the level of the cell, and promotes the accumulation of deleterious mutations (Fig. 2). However, the effect of increasing *M* remains, on an absolute scale, very mild if leakage is completely absent (Fig. 2).

**Figure 2.**
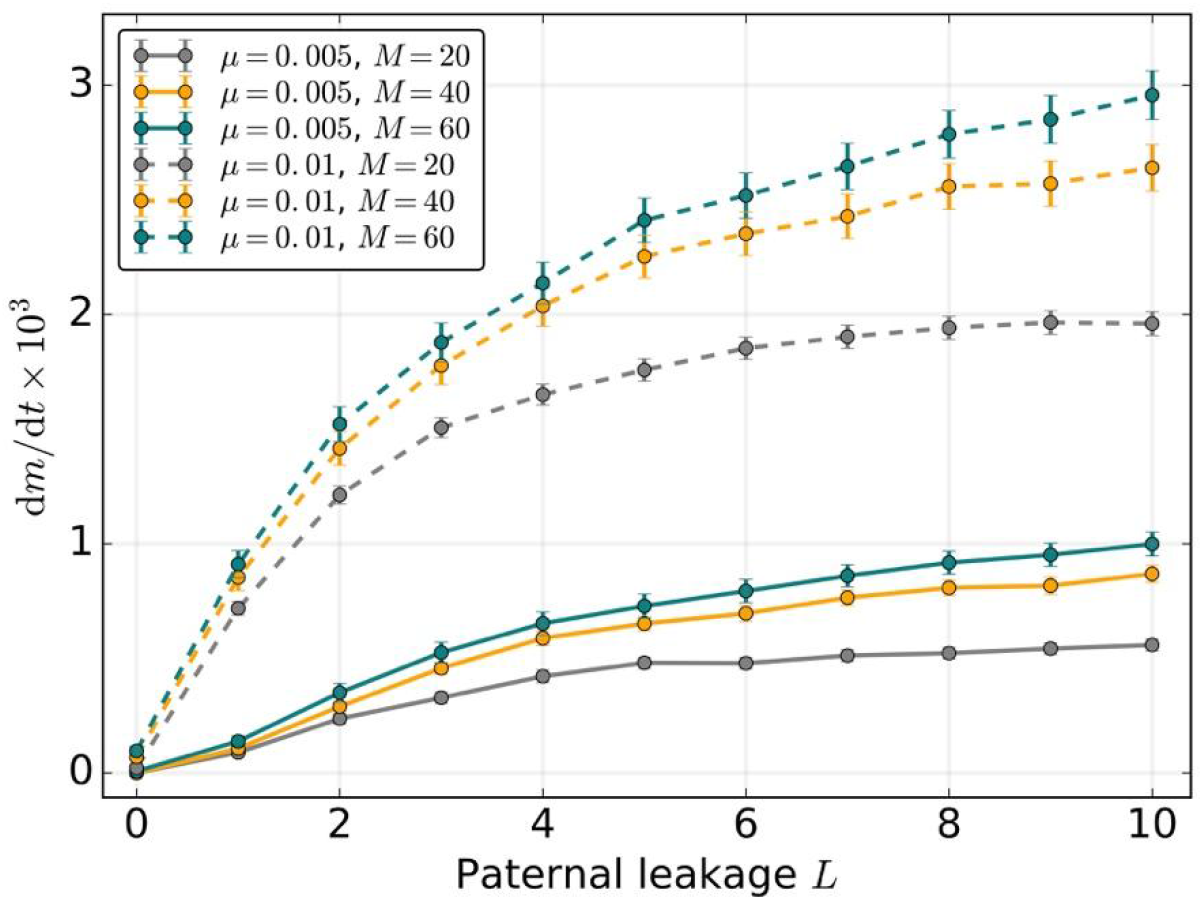
Strict uniparental inheritance of small freely-segregating mitochondrial populations mitigates the mutational meltdown in the absence of mitochondrial recombination by slowing down the rate of mutation fixation d*m*/d*t*. Mutation rate is *μ*=0.005 (solid lines) or *μ*=0.01 (dashed lines), population size *N*=500, *C*=1. Here and in the rest of the paper, error bars indicate the 95% confidence intervals for the standard error of means 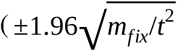, where *m*_fix_ is the number of fixed mutations after *t* generations.)

The above results assume that segregation of mitochondrial genomes is random at cell division (*C*=1). Because clustering of mitochondrial DNA molecules into linked groups such as nucleoids or organelles could curtail the variance-increasing capacity of segregational drift (Raap et al., 2012), we next investigated the effects of limited segregation by varying the size of the genome cluster *C*, and the rate of migration between clusters of the same cell *F*.

In the limit of *C*=1, all *M* copies of the mitochondrial genome segregate independently, while with *C*=*M* all *M* copies are transmitted together, each cell division producing a daughter cell identical to the parent. With genome clusters of permanent composition (i.e. no inter-cluster migration, *F*=0), weakened segregation and less efficient purifying selection leads to fast accumulation of mutant alleles (Fig. 3). However, our results show that even low rates of genome migration between clusters are capable of restoring the beneficial stochastic effect of segregational drift, sufficient to virtually eliminate the mutational ratchet-like genome deterioration (Fig. 3).

**Figure 3.**
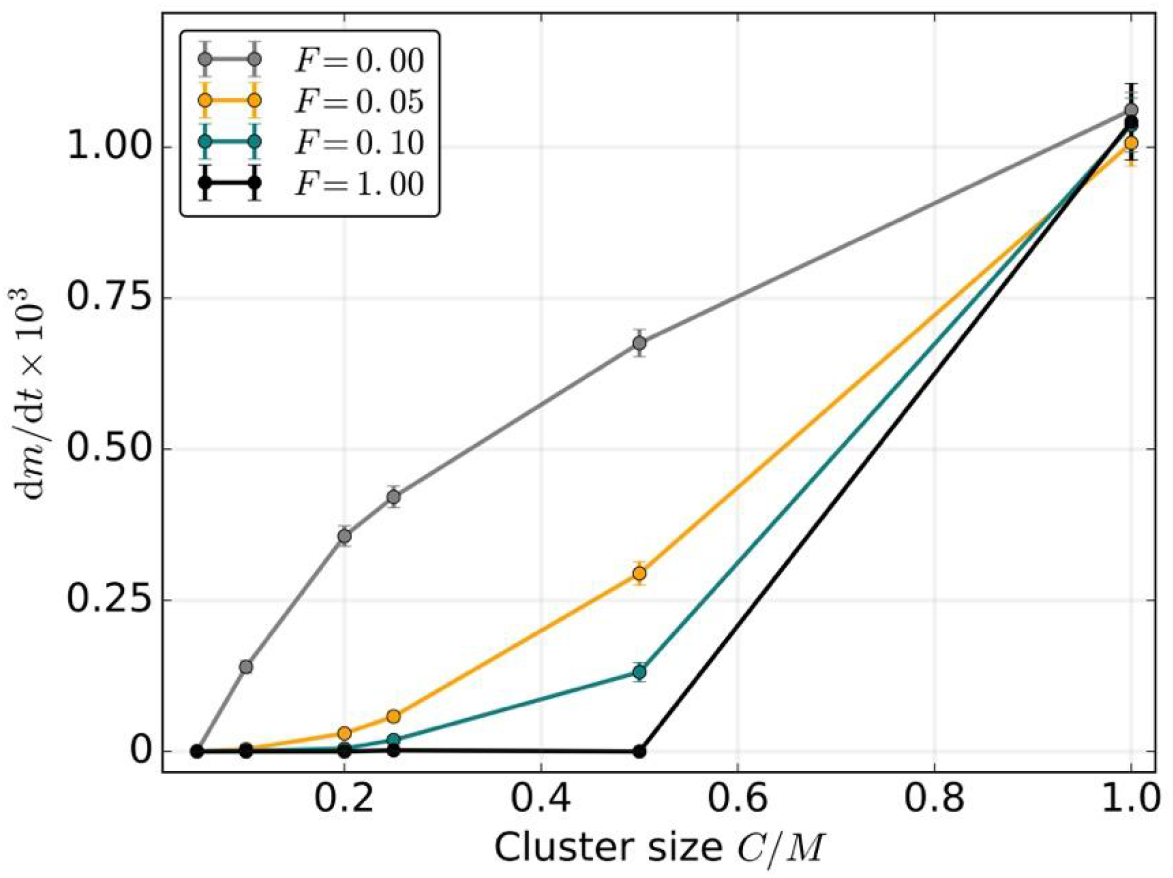
Non-random clustering of mitochondrial genomes into tightly linked groups of size *C* promotes the accumulation of weakly deleterious mutations due to suppression of segregational drift at cell division. However, if the clusters are allowed to exchange genomes between cell divisions, even very low migration rates *F* prevent the operation of Muller’s ratchet. *F* is the number of genome migration events per cell per generation, *μ*=0.005, *N*=500, *M*=20, *L*=0.

Mitochondrial populations of eukaryotic cells undergo constant transformation that involves fusion into dynamic networks, allowing the exchange of proteins, lipids and DNA, and fission, producing new organelles that differ in their protein and genomic contents (Westermann, 2010). This dynamic behavior has been suggested to serve as a mechanism of mitochondrial quality control through differential segregation of damaged mtDNA molecules and through mitochondrial autophagy (Twig et al., 2008; Kowald and Kirkwood, 2011; Hoitzing et al., 2015). Our results indicate, however, that even without selective segregation or removal of damaged genomes, high efficacy of cell-level selection can be achieved through random redistribution of mitochondrial genomes to new individual organelles, slowing down the mutational erosion of asexual mitochondrial genomes.

### Recombination slows down the ratchet, but does not reduce the equilibrium mutation load

An oft-cited advantage of meiotic sex in eukaryotes and horizontal gene transfer in prokaryotes is that recombination can restore the least-loaded nuclear genome class if the mutant allele is not fixed (Muller, 1964; Felsenstein, 1974; Takeuchi et al., 2014). To better understand the role of recombination in protecting organelle genomes from Muller’s ratchet in the more complex case of multi-copy mitochondrial genetics with purifying host-level selection, we further investigated the combined effects of paternal leakage and recombination.

In mitochondrial populations of modern eukaryotes, the scope of recombination is limited by genetic homogeneity within the cell, which is itself a result of segregational drift with uniparental transmission. Under low rates of paternal leakage, homologous recombination remains ineffective as it operates among chromosomes of largely identical composition. For low levels of mitochondrial mixing *L*, the rate of mutation fixation therefore depends only weakly on recombination rate *R* (Fig. 4A, B). With increasing rates of paternal leakage *L* and low *R*, deleterious mutations accumulate faster due to reduced strength of selection at the level of the cell (Fig. 4).

**Figure 4.**
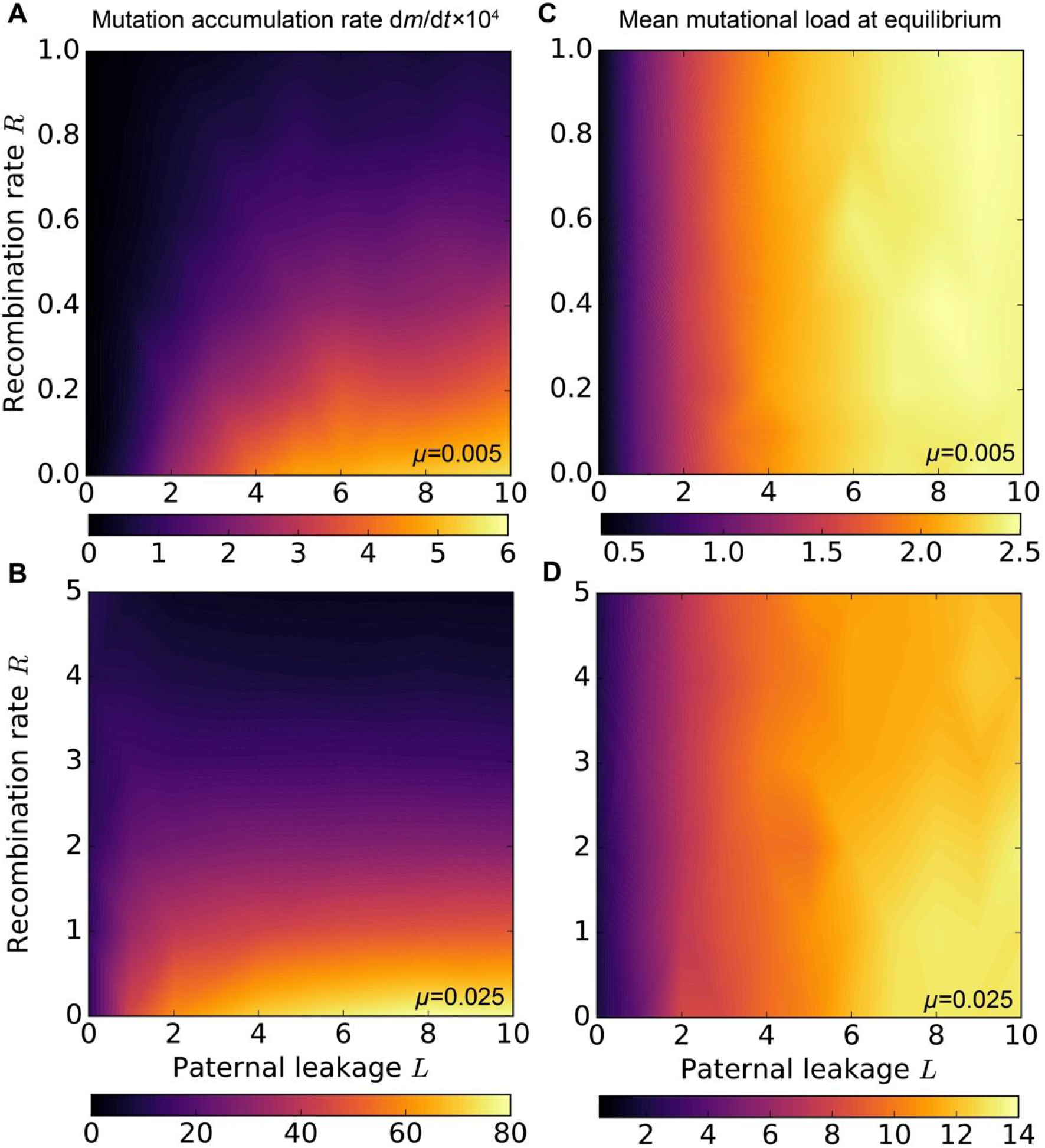
Homologous recombination slows down the accumulation of weakly deleterious mitochondrial mutations, but requires paternal leakage, which itself—in the absence of recombination—promotes mutational erosion (A, B). With high mutation and recombination rates, paternal leakage can reduce the rate of mutation accumulation relative to uniparental inheritance (B, dark regions). Nevertheless, mitochondrial mixing in the form of paternal leakage *L* increases the mean mutational load at equilibrium, i.e. in the absence of Muller’s ratchet, regardless of mitochondrial recombination (C, D). Parameter values are *μ*=0.005 (A, C) or *μ*=0.025 (B, D), population size *N*=500 (A, B) and *N*=10,000 (C, D), *M*=20. The number of mutation in least-loaded genome class is *m*_LLC_=0 in (C, D).

Recombination becomes a better defence against Muller’s ratchet when there is significant paternal leakage resulting in heteroplasmy. Under these conditions homologous gene transfer between mitochondrial genomes within the same cell will have more opportunity to regenerate the extinct least-loaded genome classes, and so the relative effect of increasing *R* becomes more significant for high *L*. The results show that with strong paternal leakage, even relatively low levels of recombination can slow down the rate of mutation accumulation, restoring the d*m*/d*t* values characteristic of much lower levels of leakage or strict UPI (Fig. 4A). Under high mutation rates, where Muller’s ratchet operates even under complete uniparental transmission (*L*=0), high levels of paternal leakage combined with frequent gene transfer reduces the rate of mutation fixation below the levels observed under strict UPI (Fig. 4B).

The rate of Muller’s ratchet, however, is an incomplete measure of all the consequences of recombination and leakage. Although recombination slows down the operation of the ratchet if there is leakage, this is expected to co-occur with reduced inter-cellular variance in mutation load (caused by leakage), and less effective purifying selection relative to strict UPI. We measured the net effect as the mean mutational load in the steady state between stochastic mutation fixation events (i.e. with a constant value of *m*_LLC_) in larger populations of 10,000 cells (Fig. 4C, D). Our simulations show that paternal leakage always leads to higher steady-state mutant load, regardless of the intracellular gene transfer rate *R* (Fig. 4C, D). Recombination can generate mitotypes that carry fewer mutations, but cell-to-cell variance remains low due to paternal leakage. As a result, deleterious mutations are not segregated out of their cytoplasmic backgrounds and are not getting purged as efficiently as they are under UPI.

### Mitochondrial bottleneck as an alternative strategy redistributing mutational variance

The mitochondrial bottleneck is an alternative strategy of redistributing mutational variance and regulating the strength of selection within and between groups of mitochondria, in many ways analogous to random genome segregation in uniparental mitochondrial transmission (Roze et al., 2005, Johnston et al., 2015). As suggested before (Bergstrom and Pritchard, 1998; Christie and Beekman, 2017), we find that tight bottlenecks reduce the long-term genome deterioration due to Muller’s ratchet, mitigating the mutational meltdown even with paternal leakage and in the absence of recombination (Fig. 5). Bottlenecks generally have weaker effect under strict uniparental inheritance, but becomes effective at reducing the rate of mutation fixation with high levels of paternal leakage (Fig 6A). Under relaxed bottlenecks, recombination is capable of reducing the ratchet rate down to the negligible rates typical for UPI or strong bottlenecks. At the same time, tight bottlenecking enforces higher levels of clonality within the cell (except for new mutations and paternal leakage), in which case the mutation accumulation rates become highly insensitive to homologous recombination (Fig. 6B).

**Figure 5.**
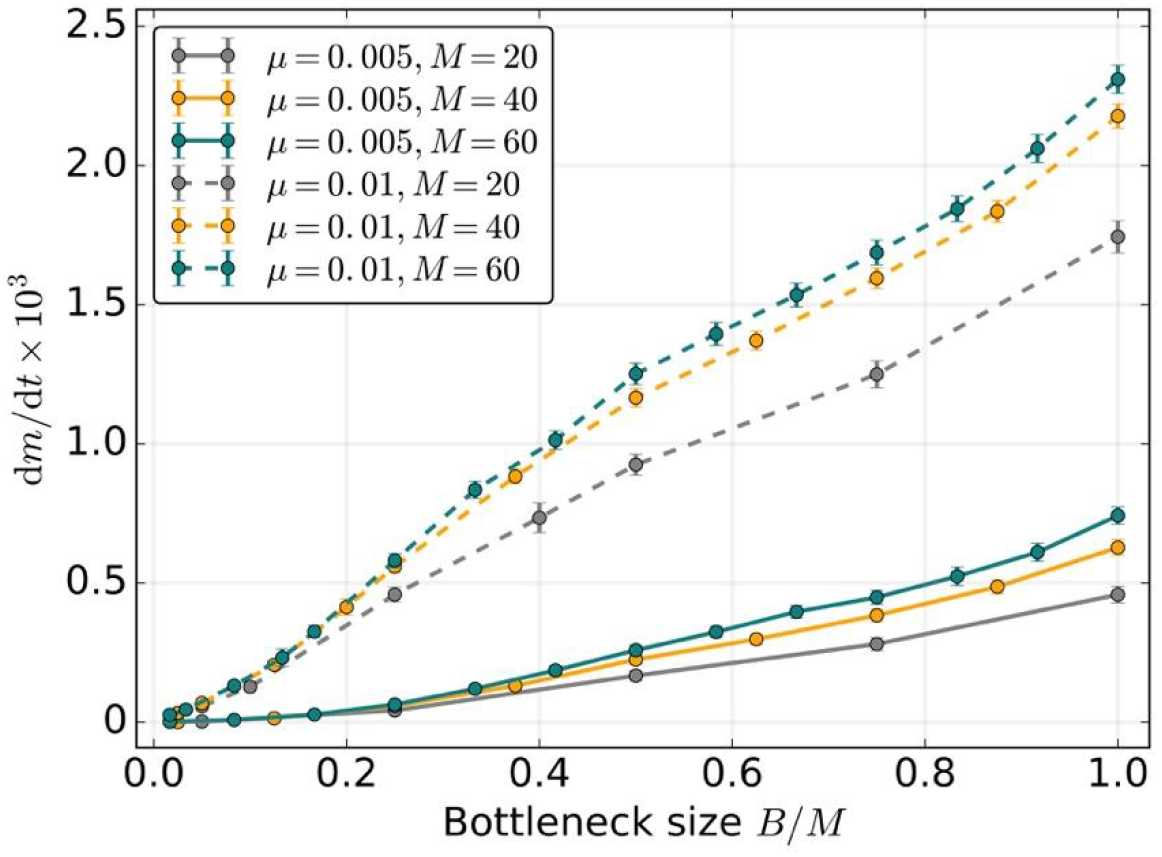
Stochastic genome resampling through mitochondrial bottlenecking reduces the rate of mutation accumulation d*m*/d*t* in the absence of recombination among mitochondrial loci. Segregational drift is less efficient in generating mutational variance among cells with larger mitochondrial populations *M*, resulting in faster rates of mutation fixation. Parameter values are: *N*=500, *C*=1, *R*=0, *L*=5.0

**Figure 6.**
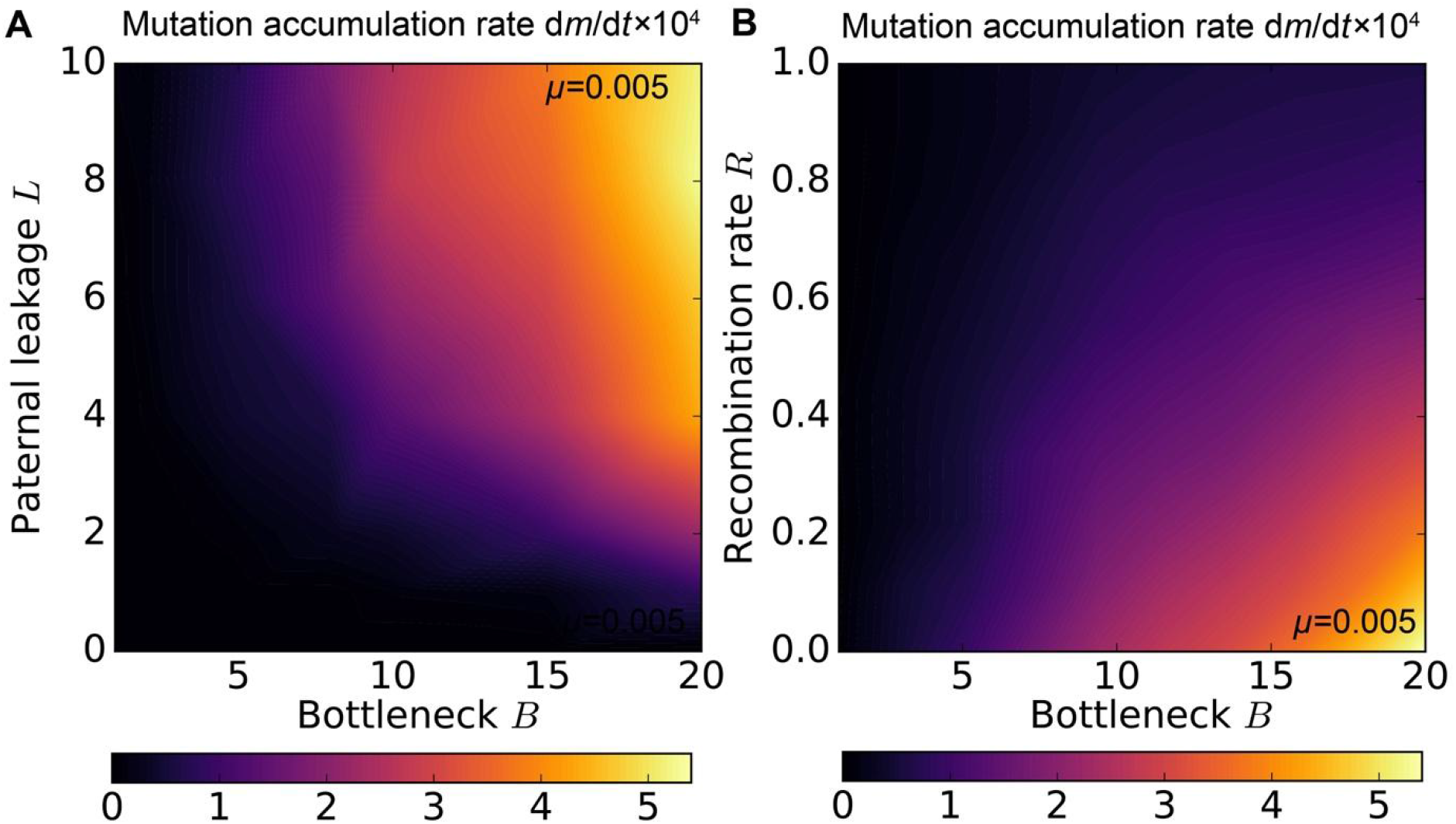
Asymmetric mitochondrial transmission and bottlenecking are two complementary strategies of increasing the cell-to-cell variance and ameliorating the mutational meltdown in the absence of recombination (*R*=0) (A). Tight mitochondrial bottlenecks increase cell-to-cell variability in mutation load and the efficacy of purifying selection at the level of mitochondrial group, and slow down the irreversible accumulation of deleterious mutant alleles. On the other hand, bottlenecks also increase clonality of mitochondrial genome within the cell, rendering homologous recombination less effective (B). *L*=5.0 (B), *μ*=0.005, *N*=500, *M*=20, *C*=1.

## Discussion

Unique features of mitochondrial population genetics evoke continuous debates over the mutational degradation of cytoplasmic organelle genomes due to the operation of Muller’s ratchet. Mechanisms that play a role in redistributing mutational variance, such as uniparental transmission and mitochondrial bottlenecks, have variously been claimed to accelerate Muller’s ratchet (Hoekstra, 2000; Neiman and Taylor, 2009; Dokianakis and Ladoukakis, 2014; Greiner et al., 2014) or to slow it down (Bergstrom and Pritchard, 1998; Roze et al. 2005; Christie and Beekman, 2017). Mitochondrial genome integrity is crucial for maintaining membrane potential and functional oxidative phosphorylation—the source of virtually all ATP of the complex eukaryotic cell. But mitochondria are predominantly transmitted uniparentally, limiting the scope and potential effects of homologous recombination.

One possible resolution of this paradox is the proposal that occasional mitochondrial recombination under biparental inheritance mitigates the mutational meltdown, and that paternal leakage could be episodically selected for (Dokianakis and Ladoukakis, 2014; Greiner et al., 2015). These proposals stem largely from early theoretical work in nuclear population genetics that established that even small levels of homologous recombination rescue finite populations from the irreversible mutational meltdown (Felsenstein, 1974; Charlesworth et al., 1993)—one of the chief evolutionary benefits of meiotic sex in eukaryotes (Kondrashov, 1993) and horizontal gene transfer in prokaryotes (Takeuchi et al., 2014).

Our study suggests that this straightforward analogy can be misleading because of important differences between mitochondrial and nuclear population genetics. Populations of mitochondrial genomes exist in a nested hierarchy of levels of selection (Rand, 2001), and are subject to segregational drift due to random partitioning at cell division, mitochondrial DNA migration within mitochondrial networks, and, possibly, germline bottlenecks in mtDNA copy numbers. While population-level drift drives the operation of Muller’s ratchet, there is also segregational drift at the level of the cell, which increases intercellular variance in mutational load, promotes host-level selection against deleterious mutations and—as our present work shows—mitigates the accumulation of weakly deleterious mutations. With low variance within the cell, homologous recombination has little effect, and is capable of restoring extinct least-loaded genome classes only in the presence of paternal leakage. But paternal leakage itself increases the equilibrium mutational load independent of population size, even if recombination rates are high.

While our current study rejects the role of paternal leakage in mitochondrial quality control, there is nevertheless a strong possibility that paternal leakage is not just a sporadic breakdown of uniparental inheritance, but is adaptive in its own right. An extraordinary array of non-conserved mechanisms that enforce the uniparental inheritance (Sato and Sato, 2013) indicates multiple origins, shifting selection pressures, and, quite possibly, reversals to partially biparental transmission of mitochondria. However, rather than being driven by the putative benefits of episodic recombination countering genome deterioration, we believe that the repeated evolution of paternal leakage is better explained by direct sex-specific selective pressures (Wade and McCauley, 2005; Kuijper et al., 2015). Note that our model examines the consequences of different rates of recombination and leakage, but does not consider how selection acts on the entity that controls these rates (e.g. nuclear genes that control the rate of paternal leakage), and this will be addressed in the future work. Sexual conflict over the control of mitochondrial inheritance provides a particularly appealing explanation for the repeated evolution of mechanisms that restrict mitochondrial transmission, frequent heteroplasmy and the prevalence of paternal leakage (Radzvilavicius, 2017). Female nuclear alleles, due to their strong linkage to the cytoplasm, favor strict uniparental inheritance, whereas male nuclear alleles, with a much weaker statistical linkage to the mitochondrial contents of the cell, would favor paternal leakage as a short-term strategy to mask detrimental mitochondrial mutations (Radzvilavicius, 2016).

Endosymbiosis at the dawn of eukaryotes produced a cell with two genomes of distinct origin, and characterized by divergent modes of inheritance and evolution. At this point, reduced selection at the lower level of individuality, e.g. selection for fittest proto-mitochondria within the cell, must have jeopardized their genomic stability due to increased deleterious mutation load or the spread of selfish genetic elements. It could be a universal feature of evolutionary transitions in individuality that mechanisms maintaining genome quality across levels of selection arise as part of the transition, and become seamlessly integrated within organism life cycles and developmental programs (Buss, 1987). For the nuclear genome of the eukaryotic cell, meiotic sex with reciprocal recombination provides one such mechanism, arising early in eukaryote evolution and, chances are, responsible for the success of the prokaryote-eukaryote transition.

But the unique population genetics of mitochondrial genes require an alternative strategy, and mechanisms increasing cell-to-cell variability in mitochondrial mutation load provide the solution. Mitochondrial fusion-fission cycles reduce linkage between mitochondrial genome copies, allowing for more efficient segregation at cell division. Stochastic genome resampling through mitochondrial bottlenecks redistributes mutational variance, increasing the efficacy of purifying selection at the level of the cell. Likewise, stochastic partitioning of mitochondria in uniparental transmission increases mutational variance between individuals, efficiently purging mitochondrial genomes that harbor excess deleterious mutations and rescuing mitochondrial genes from Muller’s ratchet in small eukaryotic populations, without the need for homologous recombination. Meiotic sex—a universal eukaryotic trait—is central to the quality control of the nuclear genome, whereas two sexes or mating types are generally required for the evolution of asymmetric organelle inheritance, and could ultimately be responsible for the long-term stability of the mitochondrial genome.

## Acknowledgements

ALR, JC and HK are funded by grants from the Academy of Finland (Centre of Excellence of Biological Interactions) and the Swiss National Science Foundation.

